# Tripartite toxin-antitoxin system regulates the production and release of pneumolysin by *Streptococcus pneumoniae*

**DOI:** 10.64898/2026.10.30.763037

**Authors:** Dora Bonini, Claire Woodall, Yuiko Takebayashi, Marcia Boura, Charlotte Richards, Murielle Baltazar, Imran Khan, Sander Granneman, Oliver J. Wilkinson, Mark S. Dillingham, Aras Kadioglu, Ruth C. Massey

## Abstract

The production of toxins is critical to the success of bacteria, with their targets and outcomes varying widely. Pneumolysin (Ply), produced by the major human pathogen *Streptococcus pneumoniae,* is a cytolytic pore forming toxin that plays many diverse roles including inducing tissue damage and inflammation, evasion of many aspects of the human immune system, invasion of the bacteria into deeper tissue, and transmission to new hosts. Despite its importance, many aspects of the biology of this toxin have to date remained unknown, in particular, what role of the three other genes that form part of the *ply* operon play in Ply production. Here we demonstrate that these genes encode a complex regulatory system, that includes a putative membrane destabilising protein (PlyA) and a MazEF-like toxin-antitoxin system (PlyB and PlyC), the toxin of which (PlyC) destabilises the Ply transcript. We propose a tripartite model whereby when the operon reaches relatively high levels of expression PlyA destabilises the *S. pneumoniae* membrane creating a stress that triggers a necessary negative feedback loop which results in the degradation of the PlyB antitoxin, release of the PlyC toxin and degradation of the *ply* transcript, thereby providing a means by which Ply production can be tightly regulated. Given the importance of Ply to the ability of *S. pneumoniae* to cause infection, identifying the factors that result in the shutdown of Ply production and release by the bacteria would seem promising target for future therapeutic strategies.

## INTRODUCTION

*Streptococcus pneumoniae* is a Gram-positive, encapsulated bacterium and the leading cause of lower respiratory infections globally [1]. This pathogen is a part of the normal flora of the human upper respiratory tract but can cause disease when it invades other body sites [2–4]. *S. pneumoniae* is responsible for a significant number of community-acquired infections, particularly in vulnerable populations such as the elderly, young children, and immunocompromised individuals [1, 5]. The global burden of *S. pneumoniae* disease has been positively impacted by the development of pneumococcal vaccines, which target the most common serotypes of *S. pneumoniae* [6]. However, the emergence of antibiotic-resistant strains and serotype switching poses ongoing challenges to public health [7–9]. Understanding the molecular mechanisms of *S. pneumoniae* pathogenesis is crucial for developing better therapeutic and preventive strategies.

*S. pneumoniae* employs several distinct approaches to cause disease. It has a polysaccharide capsule that protects the bacteria from several aspect of the host immune system [10–13]. Other surface exposed features include choline-binding proteins and pneumococcal surface protein A that help the organism adhere to epithelial cells and interfere with complement-mediated clearance [14–17]. The degradation of host tissue by secreted factors such as neuraminidase and hyaluronidase facilitate enhances the pathogenicity of the bacteria [18–21]. However, a primary means of causing host tissue damage is through the release of pneumolysin (Ply), which is a member of the cholesterol-dependent cytolysin family [22, 23]. Ply acts by forming pores in host cell membranes, leading to cell lysis, both pro- and anti-inflammatory activities, and immune system disruption; contributing to tissue damage, immune evasion and transmission to new hosts [24–30]. Ply is well established in several animal models as contributing to invasive disease. However, *in vitro* and *in vivo* studies have demonstrated that reduction in Ply production in *S. pneumoniae* subpopulations enhanced intracellular survival, and subsequently transcytosis across blood brain barrier during the development of meningitis [31, 32]. Therefore, control of when and how Ply is expressed across a population must be under stringent control to facilitate a successful lifestyle.

Unlike other bacterial toxins, Ply has neither a Sec signal sequence to facilitate generalised secretion across the bacterial cell envelope, nor a known specific secretory system. The only suggestion as to how the release of Ply is achieved by the bacteria is through autolysis and fratricide, a mechanism whereby a competent subpopulation lyses non-competent cells [33, 34]. However, this isn’t the sole mechanism of Ply release since autolysis deficient mutants can still secrete the toxin [35]. Localisation of Ply at the cell wall has also been shown to be required for the toxin’s release [36–38]. Little is also known about how the production of Ply is regulated, with some serotypes producing Ply in the exponential phase of growth and some not till stationary phase, and few specific transcriptional or translational regulators having been identified to date [39, 40]. Recently, however two new regulators of Ply production have been identified, the ZomB protein acquired on an integrative and conjugative element [41] and the *plyT* a small regulatory RNA molecule situated downstream of the *ply* gene [42]. Given the importance of Ply to the pathogenicity of *S. pneumoniae*, we sought here to characterise Ply regulation in greater detail, with a particular focus on the three uncharacterised gene co-transcribed within the *ply* operon [43], named here as *plyA, plyB* and *plyC*.

Following a detailed molecular study we have demonstrated that PlyB and PlyC act as a type II toxin-antitoxin (T-AT) system for the regulation of Ply production. The antitoxin, PlyB binds to and neutralises the toxin, PlyC. When released, PlyC binds and degrades the Ply transcript shutting down any further production of the Ply proteins. We propose that PlyA is a membrane disrupting protein that triggers intracellular stress when over expressed, leading to a negative feedback mechanism whereby PlyB is degraded, releasing PlyC which then destabilises the entire *ply* transcript. We propose that this T-AT system makes an attractive target for anti-virulence therapeutic development, as either increasing the degradation of PlyB or enhancing the activity of PlyC will result in a shutdown production and release of Ply from the bacterial cells.

## MATERIALS AND METHODS

### Strains and Growth Conditions

*S. pneumoniae* isolates were cultured on Brain-Heart Infusion (BHI) agar plates with 5% (v/v) defibrinated horse blood supplemented with antibiotics where appropriate (2 µg/ml erythromycin; 200 µg/ml spectinomycin) and incubated in 5% CO_2_ at 37°C for 18h. Colonies from the blood agar plate were inoculated into 10 ml of pre-conditioned BHI broth and two subsequent 1:100 dilutions were created then incubated in 5% CO_2_ at 37°C for 18h. The overnight broth grown culture that reached mid-log phase was sub-cultured to the required optical density (OD_600_) for subsequent experiments. *E. coli* strains were grown in LB broth supplemented with ampicillin (100 µg/ml), kanamycin (30 µg/ml), chloramphenicol (25 µg/ml) or erythromycin (200 µg/ml) where appropriate.

**Table 1:** Strains used in this study.

| Strain | Reference |
|---|---|
| *E. coli* |  |
| DH5α | Cloning strain |
| BL21(DE3) | Expression strain |
| Lemo21 (DE3) | [44] |
| ***S. pneumoniae*** |  |
| R6 | Unencapsulated D39 derivative [45] |
| R6 Δ*plyA* | This study |
| R6 Δ*plyB* | This study |
| R6 Δ*plyC* | This study |
| R6 Δ*ply* | This study |
| R6 Δ*plyB* p | This study, R6 *plyB* mutant carrying pMSP7517 empty plasmid |
| R6 Δ*plyB* p-*plyB* | This study, R6 *plyB* mutant carrying pMSP7517-*plyB* complementation plasmid |
| R6 Δ*plyB* Δ*plyC* | This study, R6 *plyB* and *plyC* double mutant |
| D39 | [46] |
| D39 Δ*plyA* | This study |
| D39 Δ*plyB* | This study |
| D39 Δ*plyC* | This study |
| D39 Δ*ply* | This study |
| D39 Δ*plyB* Δ*plyC* | This study, D39 *plyB* and *plyC* double mutant |

**Table 2:**
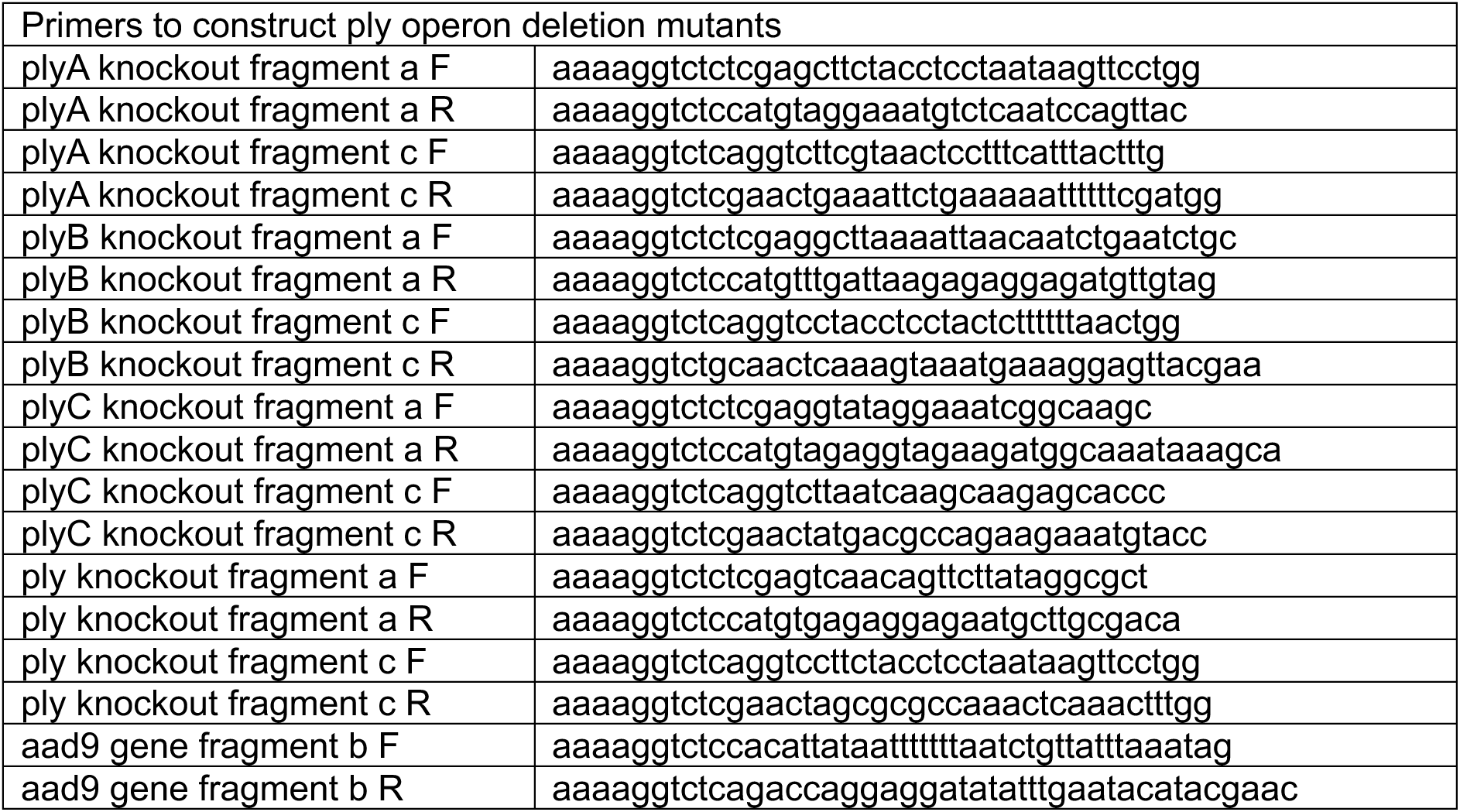
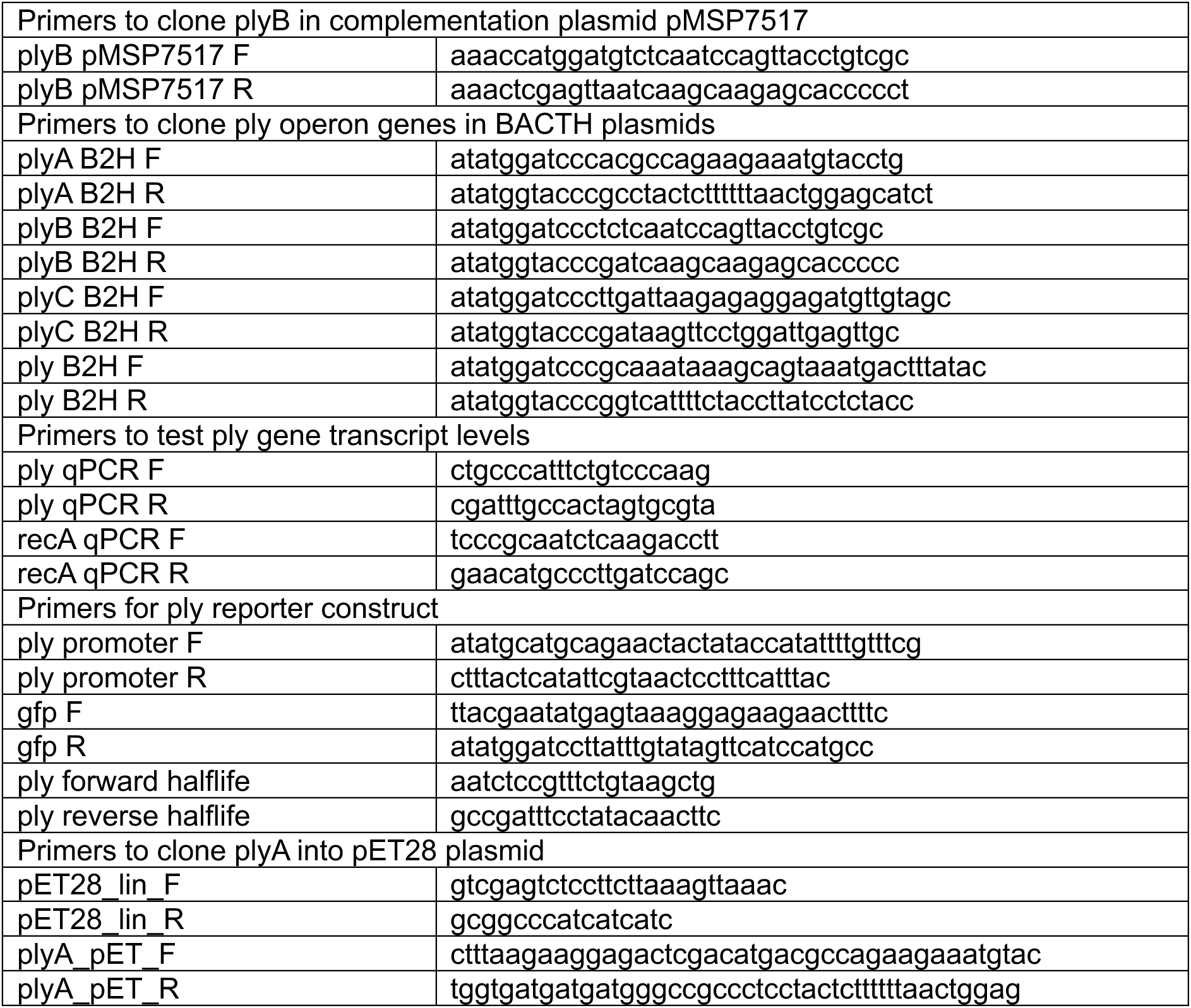
Oligonucleotide primers.

**Table 3:** Plasmids used in this study.

| Plasmid | Reference |
|---|---|
| pET28a (+) | Addgene |
| pET28a - *plyA* | This study |
| pMSP7517 | [47] |
| pVA838 | [48] |
| pCR8 | Invitrogen |
| pFW5 | [49] |
| pCN57 | [50] |
| pMSP7517 - *plyB* | This study |
| pVA838 *ply-gfp* | This study |
| pET22b – strepII- *plyB* | This study |
| pET22b – strepII- *plyC* | This study |

### Construction of deletion mutants

Genes were deleted in *S. pneumoniae* R6 using linear PCR products created by the Golden Gate Assembly protocol [51, 52]. Briefly, three fragments were amplified and assembled: (a) a 1kb fragment downstream of the gene to be deleted, (b) a spectinomycin resistance cassette (aad9, amplified from plasmid pFW5) and (c) a 1kb fragment upstream of the gene to be deleted. Primer sequence for each construct is described in Table 2. They were assembled into a vector (pCR8, Invitrogen) modified for golden gate cloning and transformed into *S. pneumoniae* as described below. To generate the mutants in D39, gDNA from the corresponding R6 mutant was used for the transformation.

### *S. pneumoniae* transformation

To transform *S. pneumoniae*, cultures were grown as previously described and, on the day, diluted to an OD_600_ 0.1, further incubated for 1 hour in BHI competency medium containing glucose (20% w/v), BSA (4% w/v) and calcium chloride (1% w/v) until OD_600_ 0.2, then diluted 1:10. To 0.2ml of the diluted cells, 5 µl of Competence-Stimulated Peptide-1 (CSP 1) (100ng/ml) was added and the cells were incubated at 37°C for 10 minutes following which 1 µg of DNA was added before further incubation at 37°C for 1 hour. A 100 ml aliquot of culture was spread onto the surface of a blood plate containing spectinomycin (200 µg/ml) and incubated in 5% CO2 at 37°C for 18h or until colonies appeared. Transformants were verified by PCR and sequencing.

### Haemolytic activity assay

The haemolytic activity of the mutants compared to the wild-type strain was determined as described previously [37]. In brief, *S. pneumoniae* strains were grown to an OD_600_ of 0.4. Samples of culture supernatant were added to 75 μl of bovine serum albumin (BSA) buffer (50 mg of BSA and 77 mg DTT dissolved in 50ml PBS) in a 96-well round bottom plate and two 4-fold serial dilutions were performed in BSA buffer. Then, 10% defibrinated sheep’s blood in PBS was washed three times and diluted to 2%. An aliquot of 50 μl 2% sheep’s blood was added to the sample serial dilutions in the 96-well plate. The plate was incubated in 5% CO_2_ at 37°C for ∼20 minutes or until the supernatant of the first dilution turned yellow and the supernatant of the second dilution stayed pink/red. The plate was centrifuged at 500 rpm for 10 minutes, then 100 μl of supernatant was transferred to a flat bottom 96-well plate and absorbance at 415 nm was recorded with a FLUOstar Omega microplate reader. Haemolysis was expressed by the haemolytic ratio (HR), where HR = (A_415_ (test) – A_415_ (negative control))/(A_415_ (positive control) – A_415_ (negative control)).

### Ply Western blots

Whole cells lysates, supernatant, cell wall and protoplasts were fractionated using modifying a previous protocol [37]. In brief, 5 ml taken from a 10 ml broth culture at OD_600_ 0.4 was centrifuged at 4,000 x g for 5 min. The pellet was saved for whole cell lysates whereas the supernatant was concentrated on a Pierce™ Protein Concentrators PES, molecular weight cut off 100 kDa, using centrifugation at 2,000 rpm for ∼45 minutes or until the supernatant had concentrated to a volume of 100 µl. For whole cell lysate preparations, the cell pellet was resuspended in 500 ml of lysis buffer and incubated for 30 minutes at 37°C and then stored at −20°C until required. To detect the Ply protein in the samples, proteins were separated on 4-12% nUView Tris-Glycine Precast gels. Western blotting was conducted using an anti-pneumolysin antibody (Abcam, catalogue number: ab71811), followed by goat anti-mouse IgG horseradish peroxidase (HRP) as the secondary The HRP signal was detected using the Metal Enhanced DAB Substrate Kit (Thermo Fisher Scientific). Band intensity was quantified using ImageJ.

### PlyA cloning and toxicity analysis

The complete *plyA* locus was cloned from *S. pneumoniae* D39 into a low copy-number pET28a(+) amplifying the plasmid with primers pET28_lin_F and pET28_lin_R and the plyA insert with primers plyA_pET_F and plyA_pET_R. This plasmid was then shuttled into the membrane protein expression strain Lemo21(DE3) for toxicity analysis. To assess toxicity, cells were grown overnight at 37°C in LB medium with selection. The next day, cultures were diluted 1:50 into 25 ml LB with selection and grown for 1.5 hours. Cultures were then normalised to an OD_600_ of 0.1 and serially diluted ten-fold in LB to a factor of 10^−5^. Dilutions were plated as 10 μl spots on LB agar containing IPTG (0 mM, 0.05 mM or 0.5 mM) and kanamycin selection. Plates were dried and incubated overnight at 37°C and photographed the following day.

### Bacterial two hybrid experiment

A bacterial two-hybrid assay was used to test the interaction between two proteins of interest in *E. coli* following the Euromedex protocol. In brief, *plyA*, *plyB, plyC* and *ply* were cloned into the *BamH*I and *Kpn*I sites of the 4 BACTH vectors, pKT25, pKNT25, pUT18 and pUT18C and sequenced to confirmed in-frame cloning. To test interactions between proteins of interest two plasmids were co-transformed in BTH101 strain which does not have endogenous adenylate cyclase activity. Three colonies per co-transformation plate were grown in LB with ampicillin, kanamycin and IPTG (1mM) for 18 hours at 30°C at 180 rpm and a 10 μl aliquot of these cultures were spotted onto LB agar plates containing ampicillin, kanamycin, X-gal (50 µg/ml) and IPTG, incubated for 24 hours at 30°C and imaged.

### Construction of *plyB* complementation strains

The *plyB* gene was amplified by PCR from R6 gDNA with primers plyB pMSP7517 F and plyB pMSP7517 R and cloned into the nisin inducible plasmid pMSP7517 at the *Nco*I and *Xho*I restriction sites to make pMSP7517-*plyB*. This plasmid and the empty pMSP7517 were transformed into the Δ*plyB* mutant as described above. An overnight culture was diluted to an OD_600_ 0.1, then exposed to nisin (100 ng/ml) and incubated for a further 3 hours or until the OD_600_ was 0.4. Cells were harvested for quantification of Ply in whole cell lysates by Western blots.

### Construction of *ply* reporter construct

The ply operon promoter was amplified from R6 gDNA with primers ply promoter F and ply promoter R. The *gfp mut2* gene was amplified from plasmid pCN57 with primers gfpF and gfpR. These two PCR fragments were combined in a third PCR reaction with primers ply promoter F and gfp R and the obtained fragment was cut with *BamH*I and *Sph*I restriction enzymes and ligated with plasmid pVA838 cut with the same restriction enzymes. The obtained plasmid was transformed in R6 and R6 Δ*plyB* as described above.

### mRNA extraction and qRT-PCR analysis

RNA was extracted from cultures grown to exponential phase. The samples were mixed with RNAprotect (QIAGEN), incubated at room temperature for 10 minutes, centrifuged at 4000 rpm for 10 minutes and the pellet stored at −70 °C. Pellets were defrosted on ice, lysed in a bead beater, with 2 cycles at 4000 rpm for 60 seconds each, with a 5-minute ice incubation between each cycle. RNA was extracted and purified using the Quick-RNA Fungal/ Bacterial Miniprep Kit, according to manufacturer’s instructions. RNA was digested with DNAseI Turbo (ThermoFisher Scientific) and 1 µg was retro transcribed to cDNA using the QScript cDNA synthesis kit (QuantaBio), according to the manufacturer’s instruction. Cycling conditions were set as follows: 1 cycle of 25 °C for 5 minutes; 42 °C for 30 minutes, 85 °C for 5 minutes, and stored at 4°C. For each reaction a negative control without reverse transcriptase was included to check for genomic DNA contamination, and the cDNA diluted to 1:50 with RT-qPCR grade water and stored at −20 °C. qRT-PCR reactions were set up in triplicate for the cDNA sample and reverse transcriptase negative control, in a 20 µL reaction using 10 µL of KAPA SYBR Fast qPCR mastermix, 0.4 µL of 50X KAPA SYBR Fast ROX Low, 0.4 µL of forward primer, 0.4 µL of reverse primer, 2 µL of template. Ct values were obtained with the Mic qPCR cycler (Bio molecular systems) and fold-change in gene expression was calculated as 2^−(ΔCt gene of interest ‒ ΔCt housekeeping gene)^, normalised to *recA* as a housekeeping gene.

### Expression and purification of PlyB and PlyC

A synthetic gene codon-optimised for *E. coli* (Geneart, Invitrogen) encoding either N-terminally StrepII-tagged PlyB or C-terminally StrepII-tagged PlyC were individually cloned into expression vector pET22b using *Nde*I and *Xho*I restriction sites. For large scale expression, the plasmids were transformed into BL21(DE3). Liquid LB cultures were grown to an OD_600_ of 0.6 and induced overnight at 18°C with 0.5 mM IPTG. The pellets were harvested by centrifugation and lysed into buffer containing 50 mM Tris pH 8.0, 300 mM NaCl, 1 mM TCEP, 10% glycerol, 1mM PMSF and protease inhibitor cocktail (Roche), then sonicated on ice for a total of 90 seconds. After centrifugation at 4°C for 30 mins at 50000g, the cleared lysates were applied to Streptactin-XT beads (GE Healthcare) in batch and incubated for 1 h at 4°C with rotation. After washing twice in batch with buffer containing 20 mM Tris HCl pH 8.0, 300 mM NaCl, 0.5 mM TCEP, once with the same buffer containing 1M NaCl, and twice again with buffer containing 20 mM Tris HCl pH 8.0, 300 mM NaCl, 0.5 mM TCEP, the protein was then eluted in the same buffer containing 50 mM biotin. The PlyB or PlyC-containing fractions were spin concentrated (10,000 Da cutoff, Millipore) down to a volume of approximately 250 µL and applied to a Superdex S75 10/300 column in buffer containing 20 mM Tris HCl pH 8.0, 200 mM NaCl, 0.5 mM TCEP. The collected peak fractions were then concentrated using a centrifugal filter unit (Millipore), glycerol added to a final concentration of 10% (v/v) and the protein stored at −80°C.

### PlyB/C degradation and recovery assay

Total RNA from 5 ml of cultures was extracted from cultures grown to stationary phase. For the degradation assays, varying concentrations of purified PlyB or PlyC were incubated with 10 µl of DNase-treated total RNA (100 ng/µl) in 25 mM HEPES (pH 8.0), 250 mM NaCl, 0.5 mM TCEP, 10% glycerol (total volume of 20 µl) at 37°C for 30 min. For recovery assays, PlyB and PlyC were mixed and co-incubated on ice for 10 min prior to addition to total RNA. The reaction mixtures were cleaned up (RNeasy mini kit, Qiagen), eluted in 20 µl RNase-free water and converted to cDNA as described above. qPCRs were performed using the primers listed in Table 2 with PowerUp SYBR Green Master Mix (Applied Biosystems) and fold-change of target gene expression normalised to housekeeping gene *recA*.

### Ply transcript half-life quantification

An overnight culture of wildtype and *plyB* mutant was added to 20 mL of TSB broth and grown to exponential phase. At time point 0, 1.5 mL aliquot of culture was harvested and mixed with RNAprotect as described above. Rifampicin (200 μg/mL) was then added to the culture, incubated and samples collected at 5, 15, 30 and 60 minutes using the same method as for the time point 0 sample. RNA was extracted as described above and the RNA integrity verified and quantified using a Qubit4 Fluorometer. 10 µg of RNA was digested with DNAseI Turbo kit in a 100 µL reaction according to manufacturer’s instructions, to ensure removal of genomic DNA. The RNA was reverse transcribed as described above.

The reactions were set up as described above with primers targeting *ply* (Table 2). Reaction conditions were set up as 95 °C for 3 minutes, followed by 40 cycles of 95 °C for 15 seconds and 60 °C for 1 minute. A melt curve was run from 60°C to 95 °C increasing 1.6 °C per second. The mean cycle threshold (Ct) values from technical triplicates, obtained using QuantStudio Design and Analysis Software (v1.5.2), were used to calculate the relative transcript abundance for each time point relative to 0 minutes. Fold changes were calculated and one phase decay analysis was performed in prism (v10.2.0) to determine mRNA half-lives for wild type and Δ*plyB*, in biological triplicate, following the methods described in Ratnadiwakara *et al* [53]. During initial analysis, the wild type showed increased *ply* mRNA abundance at 5 minutes compared to 0 minutes, indicating a biphasic decay pattern. To account for this, decay analysis was repeated using the 5-minute time point as the reference.

## RESULTS

### PlyB contributes to the haemolytic activity of *S. pneumoniae*

The *ply* gene is situated in a highly conserved operon with three uncharacterised genes upstream of it, named here as *plyA, plyB* and *plyC* (fig. 1A). To understand what role these genes play in Ply production and activity we inactivated each gene by allelic replacement with a spectinomycin resistance cassette in the genetically tractable *S. pneumoniae* strain R6 and quantified the effect that this had on the haemolytic activity of the bacteria. Inactivation of the *ply* gene, as expected, completely abolished the lytic activity of the bacteria (Fig. 1 B&C). Inactivation of *plyA* and *plyB* also significantly reduced the haemolytic activity of the bacteria, but there was no significant effect following the inactivation of *plyC* (Fig. 1B&C). Western blotting of the bacterial supernatant and of whole cell lysates supported the haemolytic activity data for the *ply* and the *plyB* mutant in that there was no Ply detectable in the *ply* mutant supernatant, and a significant reduction in Ply abundance for the *plyB* mutant (Fig. 1 D-G). To confirm that these observations weren’t specific to the R6 background, we recapitulated the *ply* operon mutants in the D39 background and confirmed the effect of *plyB* deletion on reducing Ply production (Fig. S1). While there was a slight reduction in Ply abundance in the supernatant of the R6 *plyA* mutant (Fig. 1B&C), this was not statistically significant, and there was no effect on the haemolytic activity or Ply abundance in the D39 *plyA* mutant (Fig. S1), suggesting this minor effect was specific to the R6 background.

**Fig. 1:**
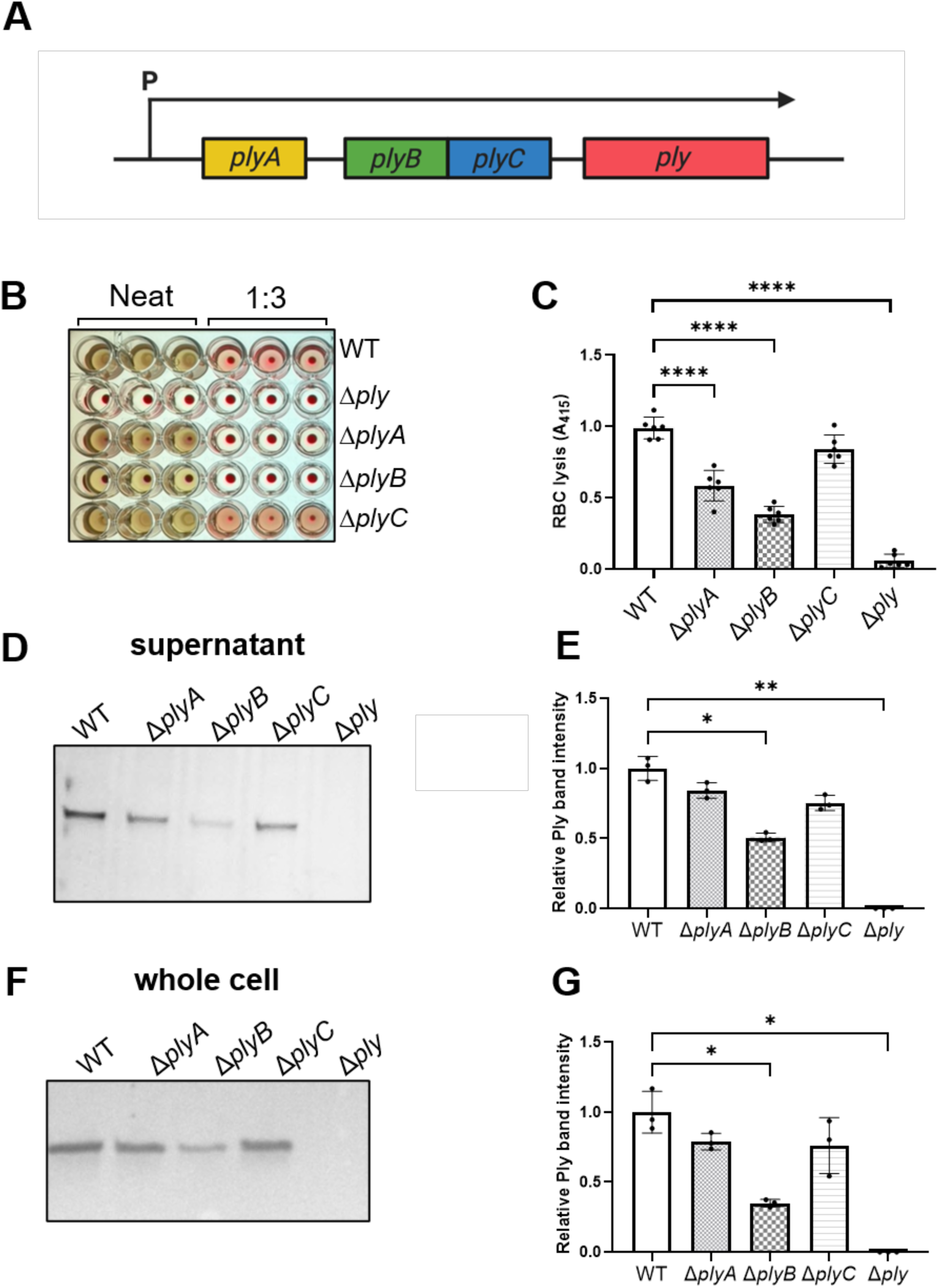
The pneumolysin operon and activity and abundance of the toxin in *plyA, plyB* and *plyC* mutants. (A) Cartoon representation of the ply operons and the three uncharacterised upstream genes *plyA, plyB* and *plyC*. (B) Representative photograph of the red blood cell lysis of the supernatant of the *ply* mutants. (C) Red blood cell lysis caused by the supernatant of the *ply* mutants (One-way ANOVA, WT vs Δ*plyA* p < 0.0001, WT vs Δ*plyB* p< 0.00001, WT vs Δ*ply* p< 0.00001). (D&E) Ply abundance in the supernatant of the *ply* mutants, (D) one representative Western blot image and (E) mean band intensity of triplicate Western blots (One-way ANOVA, WT vs Δ*plyB* p = 0.0398, WT vs Δ*ply* p *=* 0.0054). (F&G) Ply abundance in the whole cell lysates of the *ply* mutants, (F) one representative image and (G) mean band intensity of triplicate blots (One-way ANOVA, WT vs Δ*plyB* p = 0.0301, WT vs Δ*ply* p *=* 0.0163).

### Structure-function prediction analysis of PlyA, PlyB, and PlyC

Given the potential role of the other genes in the *ply* operon to affect Ply abundance, we sought to understand their activity in greater detail by searching for similar proteins based on both structural (using foldseek with Alphafold 3 (AF3)) [54, 55] and sequence (using blastp) homologies. Based on structure, the PlyA protein is predicted to be a short membrane-spanning protein with two transmembrane helices (fig. 2A). Further structural and sequence analysis revealed that PlyA is a member of the SLATT (SMODS and LOG-Smf/DprA-Associating Two Transmembrane) family of transmembrane proteins with either cell suicide or pore forming activity [56]. Further structural analysis of PlyA highlights the presence two glutamate residues situated directly in the middle of the transmembrane helices (Fig. 2B), and such polar or charged residues in the transmembrane regions of proteins are a common motif of pore forming effector proteins (*e.g.* CRISPR protein Cam1 [57], and the *E. coli* ZorO [58] and TisB proteins [59]) allowing the passage of ions across the membrane. To confirm the activity of PlyA as a membrane disrupting protein, the gene was cloned into a tightly controlled (IPTG inducible) expression vector in *E. coli*, where at low levels of expression (0.05 mM IPTG) there was no effect on the viability of the bacteria, whereas at higher levels of induction (0.5 mM IPTG) the cells were unable to grow, providing evidence that a build-up of PlyA molecules in the membrane is toxic (Fig. 2C).

**Fig. 2:**
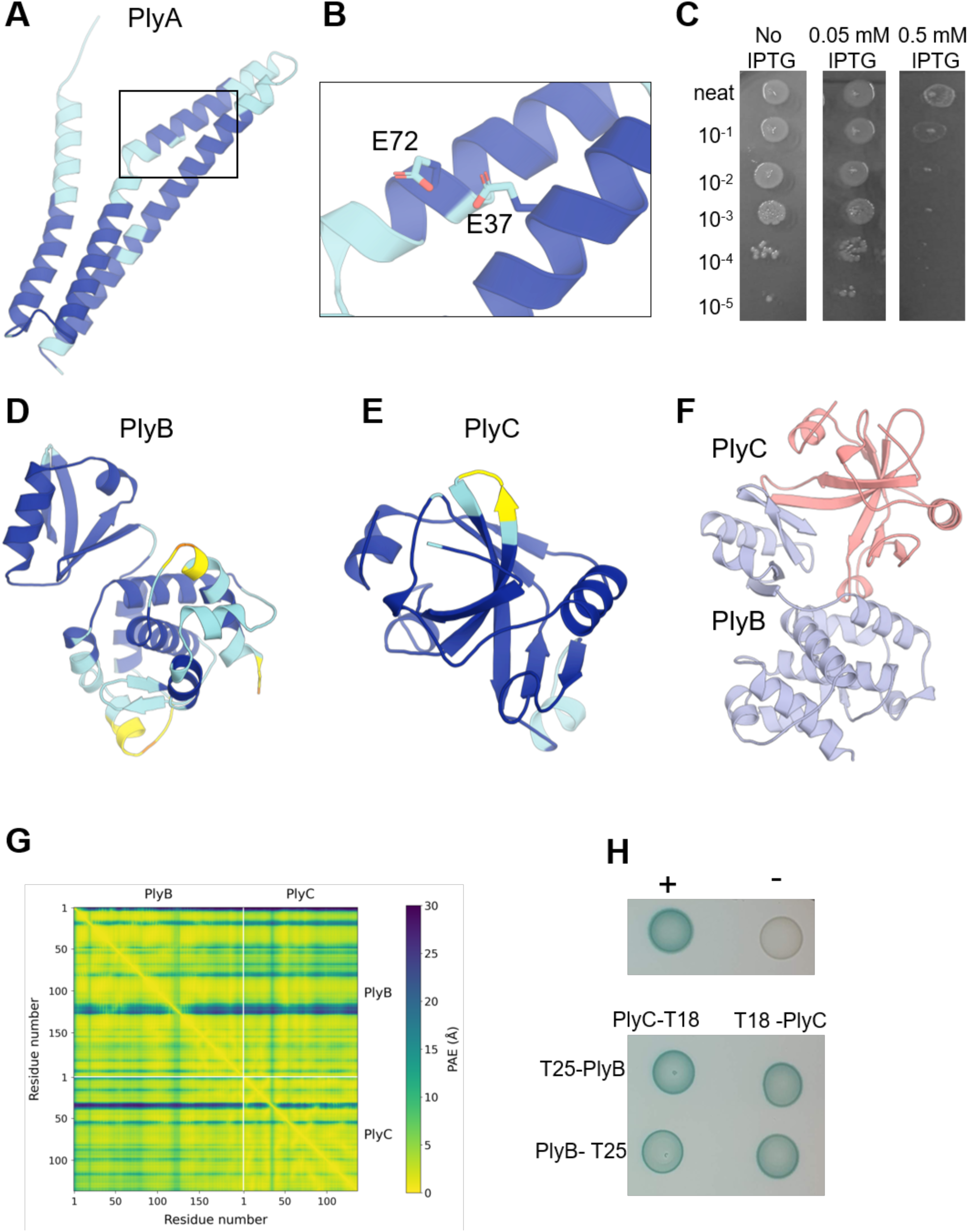
Predicted protein structures of the Ply proteins. A) AlphaFold 3 model of PlyA. The structure is coloured according to the pLDDT confidence metric (dark blue, blue, yellow and orange indicate high, moderate, low and very low confidence respectively). (B) Close up of PlyA predicted transmembrane region highlighting two glutamate residues. (C) Viability plating of Lemo21 *E. coli* cells expressing *plyA* on a plasmid under an IPTG-inducible promoter without and with increasing levels of IPTG induction. (D) AlphaFold 3 model of PlyB coloured according to the pLDDT confidence metric. (E) AlphaFold 3 model of PlyC coloured according to the pLDDT confidence metric. (F) Alphafold 3 model of the PlyBC complex (PlyB is light blue, PlyC is salmon). The protein-protein interaction interface is highly confident (iPTM = 0.91). (G) PAE scores for the PlyBC model. The relative positioning of PlyB and PlyC is highly confident (interchain PAE median = 4.1 Å; minimum = 0.87Å). (H) Bacterial two-hybrid analysis between PlyB tagged with the T25 subunit at the N-terminus and C-terminus and PlyC tagged with the T18 subunit at the N-terminus and C-terminus, as indicated. PlyB and PlyC interact. Representative of three biological replicates. pKT25 and pUT18 plasmids were co-transformed as negative control (−) and pKT25 – zip and pUT18C – zip as a positive control (+).

The PlyB protein was found to have both structural (using foldseek with AF3) and sequence (using blastp) homologies to proteins annotated as containing panacea (DUF4065) domains which are recognised as antitoxins of toxin-antitoxin (T-AT) systems [60] (Fig. 2D). T-AT systems are diverse two-gene elements widespread in bacteria where the toxin targets cellular processes and the antitoxin neutralise its effect [61, 62] Antitoxin genes are often encoded in an overlapping manner with their cognate toxin gene, and the coding regions of *plyB* and *plyC* overlap by 8 nucleotides. A search for structurally homologous proteins to PlyC identified the *Staphylococcus aureus* MazF toxin which is the toxin of a type II T-AT system, although there was little sequence homology between the proteins (Fig. 2E). Despite this, we hypothesised that PlyBC may be members of the type II T-AT family and sought to examine using AF3 whether, like other T-AT systems, PlyB and PlyC were predicted to interact. AF3 modelled a PlyB–PlyC 1:1 complex with very high confidence, with a model ranking score of 0.91, global iPTM of 0.91 and pTM of 0.92. The minimum predicted aligned error between the two chains was also low (chain-pair PAE min = 0.87 Å) (Fig. 2F&G). To experimentally validate this, we undertook a bacterial two-hybrid (BACTH) approach and found that regardless of what terminus of the PlyB and C proteins the BACTH tags (T25 and T18) were positioned, the two proteins directly interacted resulting in the production the β-galactosidase enzyme (fig. 2F). No other combination of Ply proteins was shown to interact using this methodology (Fig. S2). Together these findings suggest the PlyA, B and C may work together as novel T-AT system to control the production and release of Ply by *S. pneumoniae*.

### Characterisation of the Ply T-AT system

As a putative T-AT system, we hypothesised that inactivation of the *plyB* anti-toxin gene results in the release of the PlyC toxin, and this is what is affecting Ply abundance. We did not see an effect on Ply abundance when we inactivated the *plyC* gene alone (Fig. 1). This could be explained by PlyC being bound and inactivated by PlyB in the wild type strain in the conditions used, and as such no additional effect would be observed by deleting the *plyC* gene. As a first step we sought to complement the effect of the loss of PlyB on Ply production by cloning the *plyB* gene into the pMSP7517 expression plasmid and transforming this in the *plyB* mutant. This restored Ply production of the mutant to wild type levels (Fig. 3A&B) confirming the effect of PlyB in Ply production. Next, we sought to examine whether the effect of the loss of PlyB on Ply production was mediated via PlyC. If PlyBC is a Ply regulating T-AT system, then the deletion of *plyC* from the *plyB* mutant should restore the Ply abundance of the *plyB* mutant back to wild type levels. To test this, we constructed a double *plyBC* mutant, and quantified Ply abundance alongside the single *plyB* and *plyC* mutants by Western blotting. In accordance with its activity as a Ply abundance affecting T-AT system, the inactivation of both genes restored Ply abundance of the *plyB* mutant back to wild type levels (Fig. 3C&D). Based on these data we are confident in our hypothesis that PlyB and PlyC work together as Ply-abundance-affecting T-AT system.

**Fig. 3:**
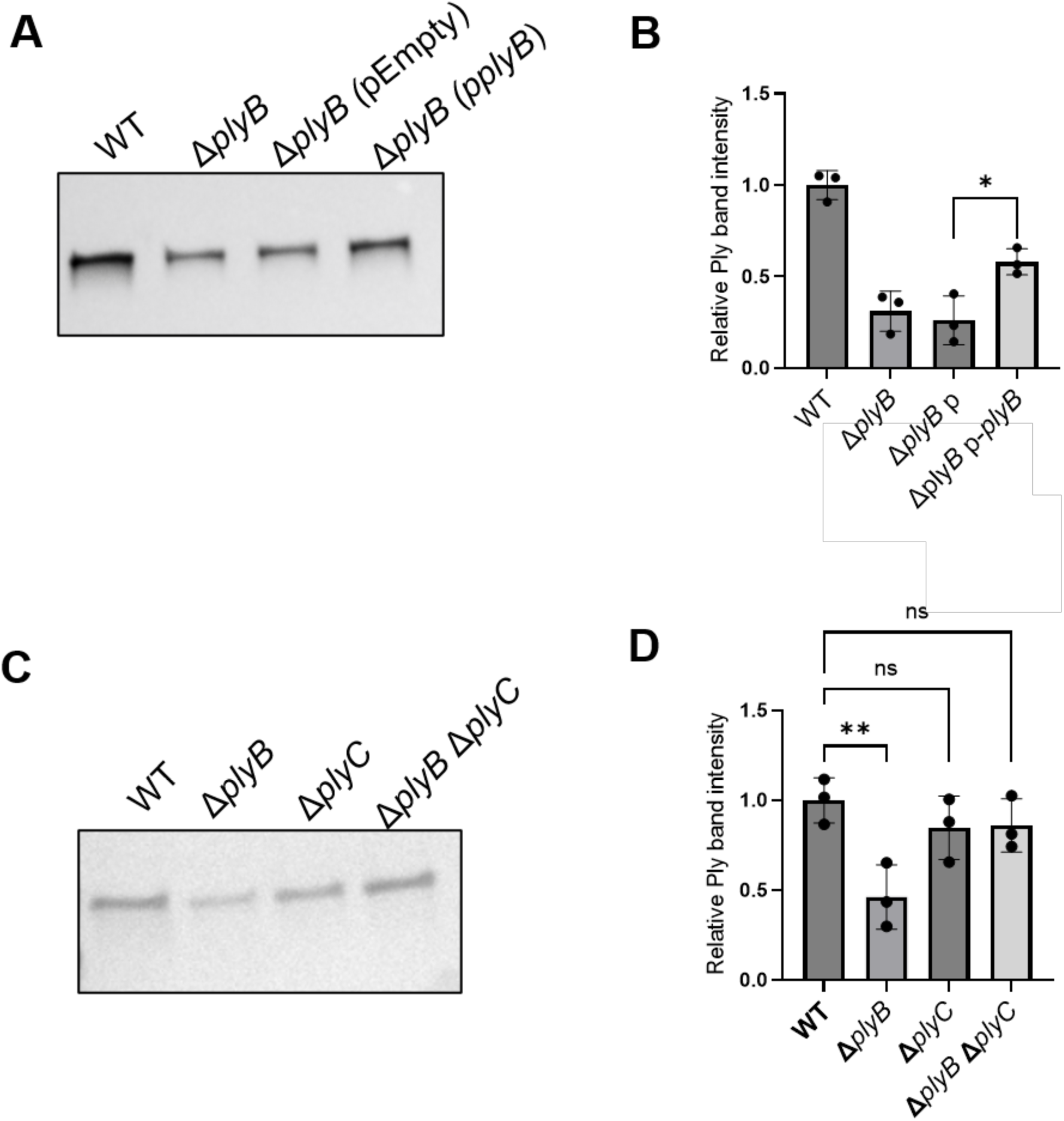
PlyB and PlyC are a Ply abundance affecting T-AT system. (A&B) Ply abundance in the supernatant of the *plyB* complemented mutant (Δ*plyB p-plyB*) compared to the *plyB* mutant carrying an empty plasmid (Δ*plyB p-*) (A) one representative Western blot image and (B) mean band intensity of triplicate Western blots (paired t-test, p = 0.0215). (C&D) The deletion of both *plyB* and *plyC* restores Ply abundance in whole cell lysate back to wild type levels: (C) one representative Western blot image and (D) mean band intensity of triplicate blots (One-way ANOVA, WT vs Δ*plyB* p = 0.0082, WT vs Δ*plyC* p *>* 0.05, Δ*plyB* Δ*plyC* p *>* 0.05).

### *In silico* predictions identify a possible PlyC RNA-binding cleft

The Ply T-AT system most resembles a type II T-AT systems, the toxins of which, such as MazF, are reported to have transcription-regulatory activity by regulating promoter activity and/or degrading transcript via ribonuclease activity. Either way, an ability to bind RNA is needed for its activity. To gain insight into the PlyC RNA-binding potential, we used the pyRBDome pipeline [63] to predict RNA-binding residues in the protein. Briefly, the pipeline employs a variety of machine learning and AI models to predict RNA-binding residues based on sequence conservation and structure (PST-PRNA [64]), presence of positively charged surface patches [65], sequence conservation alone (RNABindRPlus [66]) and whether the region is disordered [67, 68]. The pipeline then employs an XGBoost model to score each amino acid for RNA-binding propensity based on the results from the individual predictors (pyRBDome model). Representative results are shown in Fig. 4A. These analyses revealed a prominent RNA-binding cleft with a highly positively charged surface (Surface patches) and highly predicted binding propensity, with the pyRBDome consensus model providing a detailed map of the potential RNA-interaction surface. To gain structural insights into the binding mechanism, we generated an AF3 model of PlyC in complex with a single-stranded RNA sequences that is a known *B. subtilis* MazF toxin substrate ([69]; Fig. 4B). The resulting model positioned the RNA within the pyRBDome predicted cleft, with the RNA scaffold making extensive contacts across the high-probability binding surface identified by pyRBDome. These results support our hypothesis that PlyC could be a functional counterpart of MazF and act as an endonuclease, suggesting that the identified RNA-binding cleft may serve as the catalytic site for RNA cleavage. In further support of this, our model of the PlyBC complex demonstrates that PlyB binds to the same site on PlyC as the RNA binding cleft, providing a clear steric exclusion mechanism for detoxification of PlyC by PlyB.

**Fig 4:**
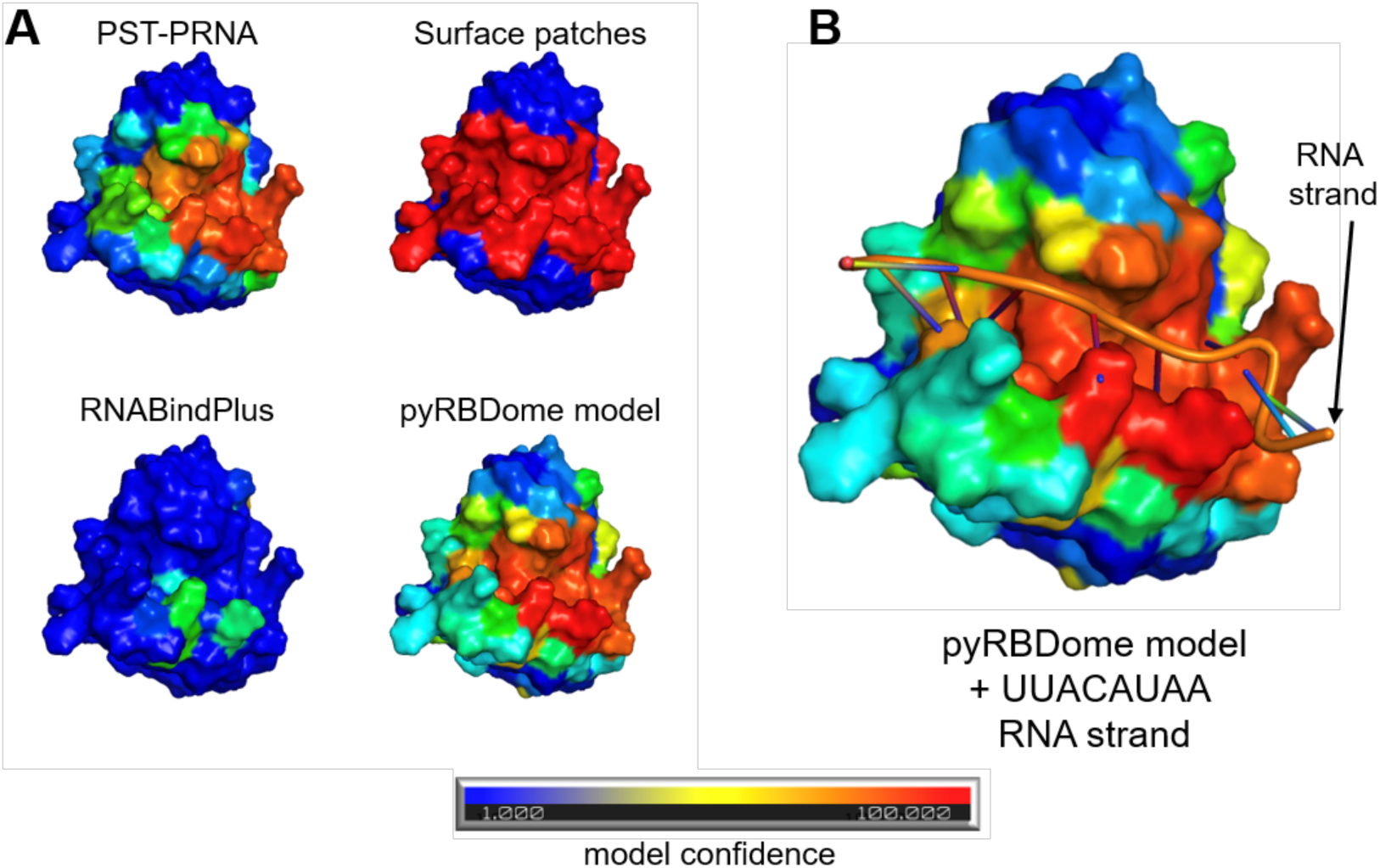
PlyC has a predicted RNA-binding cleft. (A) Comparison of RNA-binding predictions using four different methods: PST-PRNA, Surface patches detection, RNABindPlus, and pyRBDome model. Protein surface is coloured according to a spectrum gradient representing predicted RNA-binding likelihood (blue, low probability; red, high probability). Spectrum color gradient scale represents RNA-binding probability from low (blue), middle (yellow) to high (red). (B) AlphaFold 3 structural model of the PlyC in complex with a short RNA substrate (UUUACAUA; orange/brown), positioned within the RNA-binding cleft. Surface colouring reflects the same spectrum gradient, demonstrating consistency between the computational predictions and the structural model of RNA interaction.

### PlyC destabilises the *ply* transcript

To understand whether PlyC was affecting Ply abundance post-transcriptionally we tested promoter activity and *ply* transcript abundance. We first constructed a reporter plasmid with the promoter of the *ply* operon driving expression of a *gfp* gene. The plasmid was transformed into both a wild type (where PlyC is inactive) and *plyB* mutant (where PlyC is active), however no difference in GFP levels observed between these strains (Fig. 5A). We next quantified the transcript level of the *ply* gene by qRT-PCR in both the wild type and *plyB* mutant and found there to be a significant decrease in *ply* mRNA in the *plyB* mutant (Fig. 5B). Together this suggests that PlyC does not have direct transcriptional regulatory effect on *ply* promoter activity but affects mRNA levels post-transcriptionally, as might be expected for a protein with ribonuclease activity. To examine this further, we cloned, expressed and purified PlyB and PlyC proteins with a strepII tag in *E. coli* (Fig. S3). Using the purified PlyC, we extracted total mRNA from the wild type *S. pneumoniae* strain, incubated it in a range of concentrations of PlyC, and quantified the mRNA level of both the *ply* and *recA* (a housekeeping gene) gene by qRT-PCR. The *ply* transcript was degraded in a dose-dependent manner by purified PlyC (Fig. 5C). To determine whether PlyB can neutralise the activity of PlyC in this *in vitro* system, we preincubated the two proteins together before adding total RNA and determining transcript levels by qRT-PCR. While PlyB had no direct effect on *ply* transcript levels, it was able to neutralise *ply* transcript degrading activity of PlyC (Fig. 5D). We also performed a time-course experiment where transcription was halted by the addition of rifampicin and half-life of the *ply* transcript was quantified by qRT-PCR over a time-course. A decay analysis demonstrated that the half-life of the *ply* transcript in the *plyB* mutant was less than half that of the wild type strain, averaging 3.9 minutes as opposed to 11.4 minutes in the wild type (Fig. 5 E&F). Together with the structural prediction analysis, these data suggest that PlyC affects Ply abundance by destabilising and reducing the half-life of the *ply* transcript. The simplest interpretation of this is that PlyC is an mRNA nuclease specifically targeting the *ply* transcript, but this remains to be directly demonstrated.

**Fig. 5:**
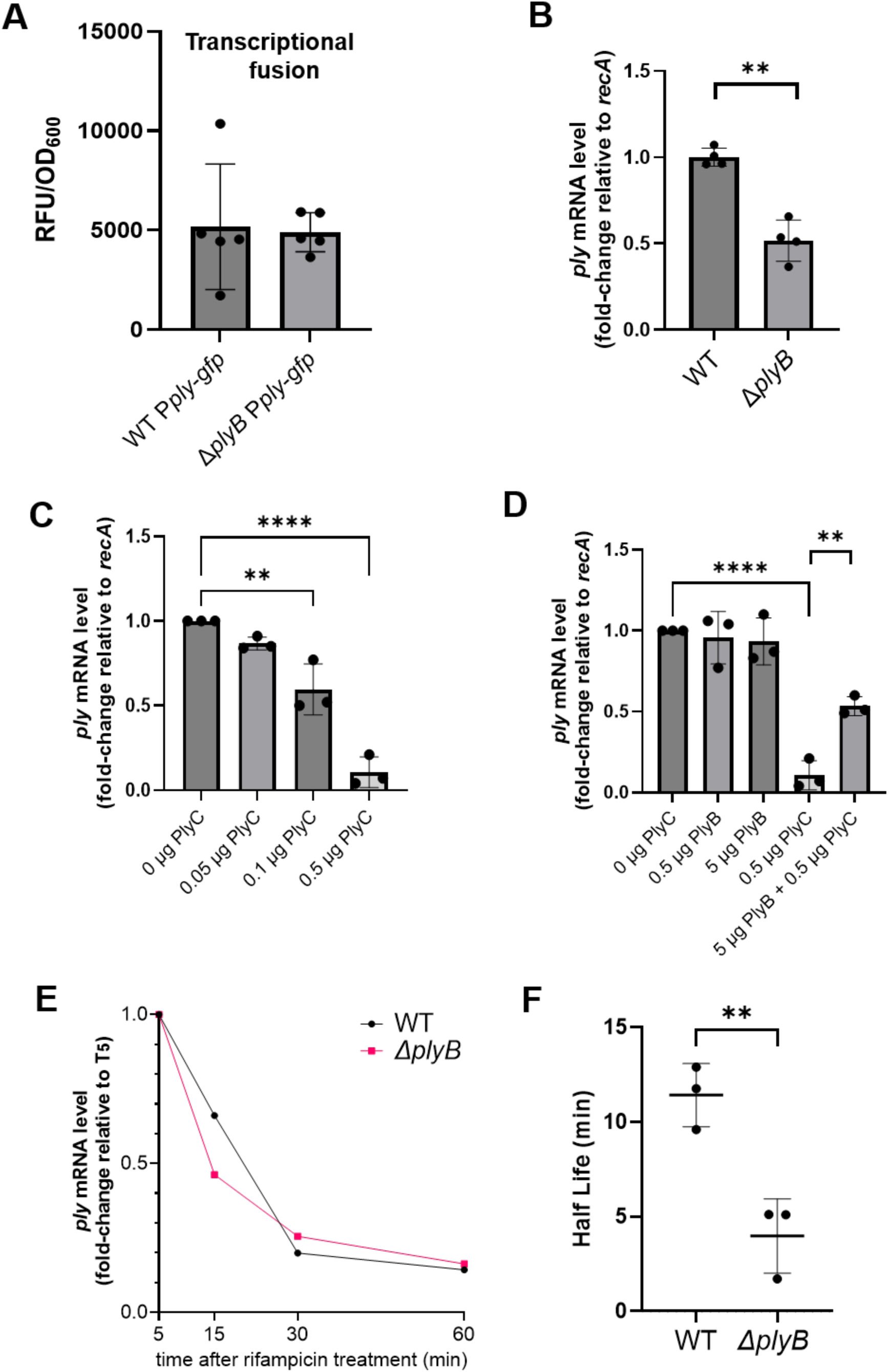
PlyC has *ply* transcript degrading/destabilising activity. (A) *ply* promoter activity in the wild type and *plyB* mutant as determined by reporter (GFP) activity shown as relative fluorescent units normalised by OD_600_. (B) Fold-change in *ply* mRNA levels relative to the housekeeping gene *recA* in the wild type and *plyB* mutant as determined by qRT-PCR (paired t-test, p = 0.0015). (C) Fold-change in *ply* mRNA levels *in-vitro* following exposure of total mRNA extracted from the wild type to increasing concentrations of PlyC, as determined by qRT-PCR normalising to the housekeeping gene *recA*. PlyC has a dose dependent effect on *ply* transcript abundance (One-way ANOVA, 0 µg PlyC vs 0.1 µg PlyC p = 0.0015, 0 µg PlyC vs 0.5 µg PlyC p= <0.0001). (D) Fold-change in *ply* mRNA levels in-vitro following exposure of total mRNA extracted from the wild type to increasing concentrations of PlyC, PlyB and PlyB and PlyC together, as determined by qRT-PCR. PlyB inactivated the endonuclease activity of PlyC. (One-way ANOVA, 0 µg PlyC vs 0.5 µg PlyC p < 0.0001, 0.5 µg PlyC vs 5 µg PlyB + 0.5 µg PlyC p = 0.0024) (E) Time course of fold-change in *ply* mRNA levels following exposure to 200 μg/mL rifampicin, as determined by qRT-PCR. (F) One phase decay analysis of the rifampicin time course assay demonstrated significantly decreased half-life of *ply* transcript in the *plyB* mutant (unpaired t-test, p = 0.0075).

## DISCUSSION

In this study we characterise the activity of the three gene encoded upstream of the Ply toxin, and in doing so propose a model whereby the *ply* operon is transcribed and translated into the four proteins PlyA, PlyB, PlyC and Ply (Fig. 6). PlyB binds and detoxifies PlyC while PlyA embeds itself into the bacterial membrane, and at this early-stage Ply remains in the bacterial cytoplasm. As transcription of the *ply* operon continues the PlyBC complex and Ply increases in abundance in the cytoplasm as does the presence of membrane destabilising PlyA in the membrane. We hypothesise that the damage to the membrane mediated by the increasing amounts of PlyA embedded there enhances the existing cell turnover activity of the bacteria, and this this has two potential roles in Ply production. The first is that it enhances the existing means by which Ply is released by causing further membrane breakdown, and the second that it induces a stress response that induces PlyB degradation and release of PlyC which then shuts down any further Ply protein production by degrading the *ply* transcript (Fig. 6).

**Fig. 6:**
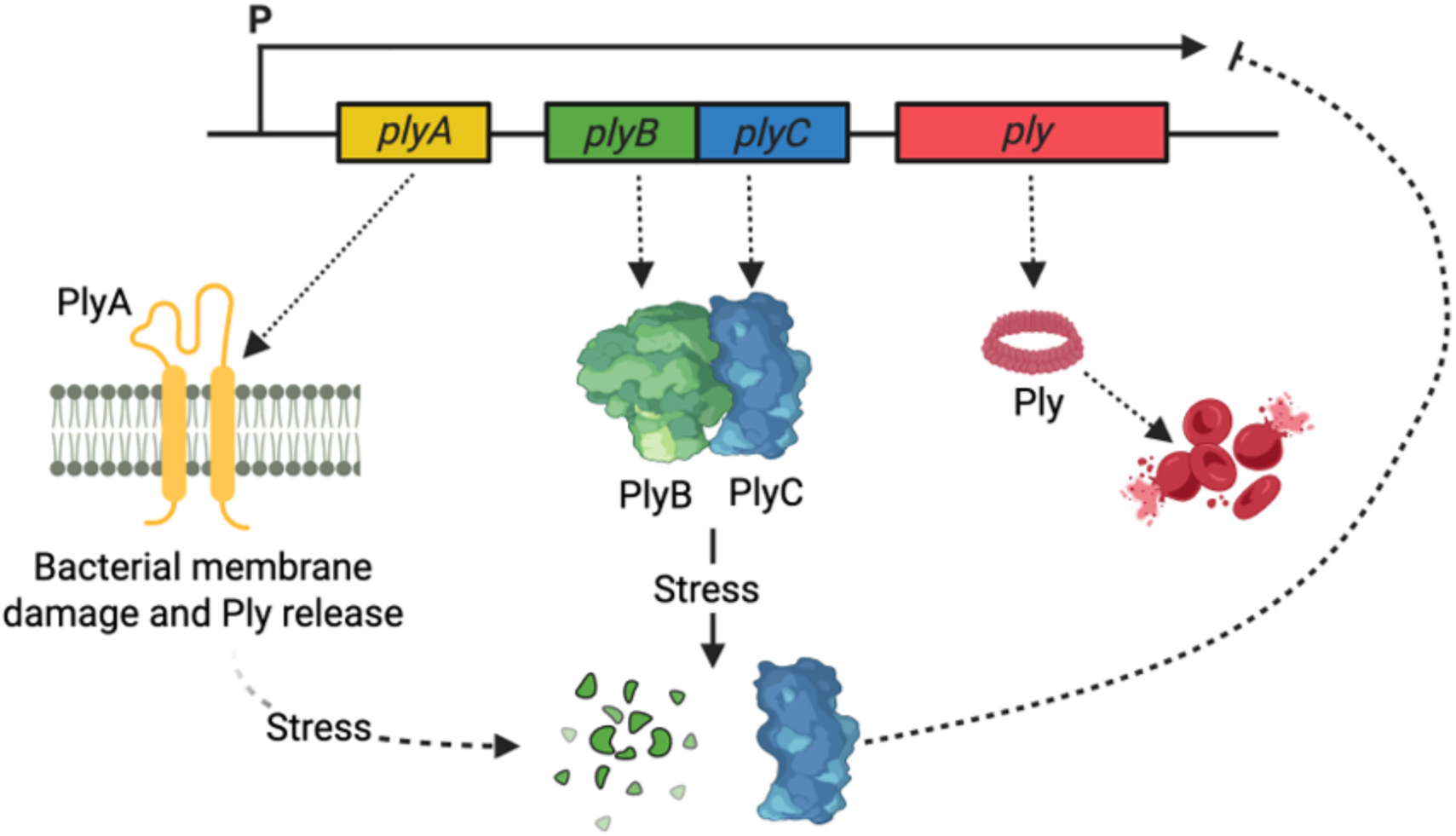
Model of proposed mechanism of regulation of Ply toxin production and release by PlyA, PlyB and PlyC. Transcription of the *ply* operon would lead to accumulation of PlyA in the bacterial membrane, with consequent damage, potentially favouring Ply release from the cell. Insertion of PlyA in the membrane could also cause generalised stress to the cell which, possibly with other stress signals, could trigger the degradation of PlyB, normally found in complex with PlyC. The degradation of PlyB causes the release of the proposed endonuclease PlyC which would in turn degrade the *ply* mRNA transcript, shutting down the production of the Ply toxin.

There are yet unproven aspects to our model. For PlyA, we only have a demonstrable effect on the haemolytic activity of one set of mutant bacteria, and a direct connection from PlyA overexpression to intracellular stress induction causing PlyB degradation is lacking. However, in *E. coli* PlyA is toxic and given the predictions of its structure and homology to SLATT proteins, its putative role in cell membrane destabilisation seems likely. Further experimentation comparing the transcriptome and/or the proteome of the wild type and the *plyA* mutant may help here, as would quantifying PlyB abundance and integrity under a range of stressful environmental condition with and without PlyA. Another limitation of this study is that we have not directly demonstrated RNA binding activity for PlyC. As such the exact biochemical activity of PlyC in relation to how it destabilised the *ply* transcript remains to be determined.

T-AT systems were originally identified associated with mobile genetic elements (MGEs) such as plasmids and were believed to act as an addiction system resulting in the preservation of the plasmid in the host bacterium. Since then, more diversity in activity has been reported with several chromosomally encoded T-AT systems having been identified, however, these have largely been associated with other type of chromosomally embedded MGEs such as phage. Despite being a naturally competent and highly recombinogenic bacterium, there has been no suggestion to date the *ply* operon is encoded on an MGE. However, a more detailed analysis if the region upstream of *ply* reveals that it is AT-rich and also contains a gene with homology to a prophage regulator (spr1745), so it is possible that this operon is within, or adjacent, to an old MGE.

It is interesting to note that had we been limited to analysing these proteins based on sequence similarity alone, we would not have identified the PlyBC proteins as being a T-AT system. While sequence analysis did identify a panacea (DUF4065) domain in PlyB, which are recognised as antitoxins of toxin-antitoxin (T-AT) systems, it was only through the use of Alphafold3 that the structural similarity between PlyC and the *Staphylococcus aureus* MazF toxin was identified, as there is little sequence homology between the proteins. In the era of great concern about the use of AI in research, here is a positive example of it helping us make a significant leap in our understanding of the fundamental biology of a bacterial pathogen.

The conservation of the Ply operon structure not only in *Streptococcus pneumoniae* strains but also other closely related Streptococcal species, *S. pseudopneumoniae* and *S. mitis* [70], suggests that it is ancient and under strong selective pressure. This is intriguing, considering how many considerably more simplistic toxin regulatory systems exist in bacteria. For a highly competent and recombinogenic bacteria, there must have been ample opportunities for the bacteria to combine the Ply gene with a more straightforward regulatory and secretory system, and yet this PlyABC system has prevailed. It is possible that there is some of the traditional addiction aspects to these genes which is why they have remained and are highly conserved. Anecdotally, we observed regular but inconsistent growth defects upon the expression of PlyC in both *S. pneumoniae* and *E. coli*, but when we tried to quantify these, which involved some necessary passaging of the bacteria, they seemed to resolve. Therefore, it is possible the PlyC has some additional toxic effect on *S. pneumoniae* that bacteria in the laboratory can compensate for upon passage, but that elicits a more detrimental effect out in their natural environment, which might explain its success.

The interesting question that this work brings up is whether it can be exploited for clinical usage. Overexpression of the *ply* operon and PlyA production might lead to an increase in lysis of the bacteria, but also an increase in expression and release of Ply, so unlikely to be of overall clinical benefit. If we instead consider post-transcriptional activity, a means of either increasing PlyB degradation and/or release of the PlyC may have greater potential. Enhanced degradation of the *ply* transcript would have immediate benefits both during infection and given the role of Ply in transmission between hosts, it could also contribute to slowing outbreaks. If as our anecdotal evidence mentioned above, increased PlyC also has an overall toxic effect on the bacteria, then this could further the clinical potential of focussing on it as a novel therapeutic target.

## Data Availability Statement

The authors confirm that the data supporting the findings of this study are available within the article and supplementary material. Bacterial strains developed as part of this study are available upon request.

## Author Contribution Statement

DB and RCM conceived the underlying idea. DB, CW, YT, MB, CR, MB, IK, OJW performed the experiments and collected and analysed the resulting data, and prepared the figures. SG and MSD undertook the structural analysis of the proteins, and prepared the figures. AK provided resources and was involved in data analysis and figure preparation. All authors contributed to the writing of the manuscript with DB and RCM responsible for final editing. RCM secured the funding for this work.

## Acknowledgements

The authors would like to thank the following people for helpful discussions as we worked our way through understanding this regulatory system: Nicholas Croucher, Henrik Strahl, Tracy Palmer, Giusy Mariano, Thomas Wood, Gareth McVickers, Gemma Atkinson, Vasili Hauryliuk, Tim van Opijnen, Marco Oggioni, and all members of Cork and Bristol Massey Group.

## SUPPLEMENTARY FIGURES

**Fig. S1.**
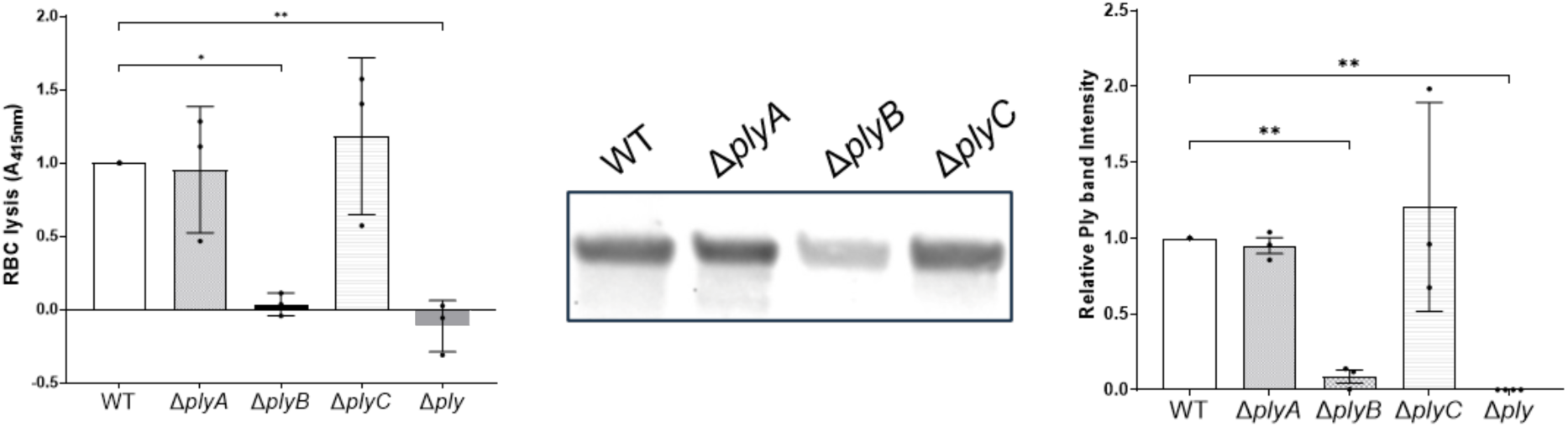
Effect of PlyA, PlyB and PlyC on Ply abundance in D39. (A) Red blood cell lysis caused by the supernatant of the *ply* mutants in the D39 background (One-way ANOVA, WT vs Δ*plyB* p = 0.0133, WT vs Δ*ply* p = 0.0054). (B&C) Ply abundance in the supernatant of the *ply* mutants in the D39 background, (D) one representative image and (E) mean band intensity of triplicate blots (One-way ANOVA, WT vs Δ*plyB* p = 0.0044, WT vs Δ*ply* p *=* 0.0012)

**Fig. S2.**
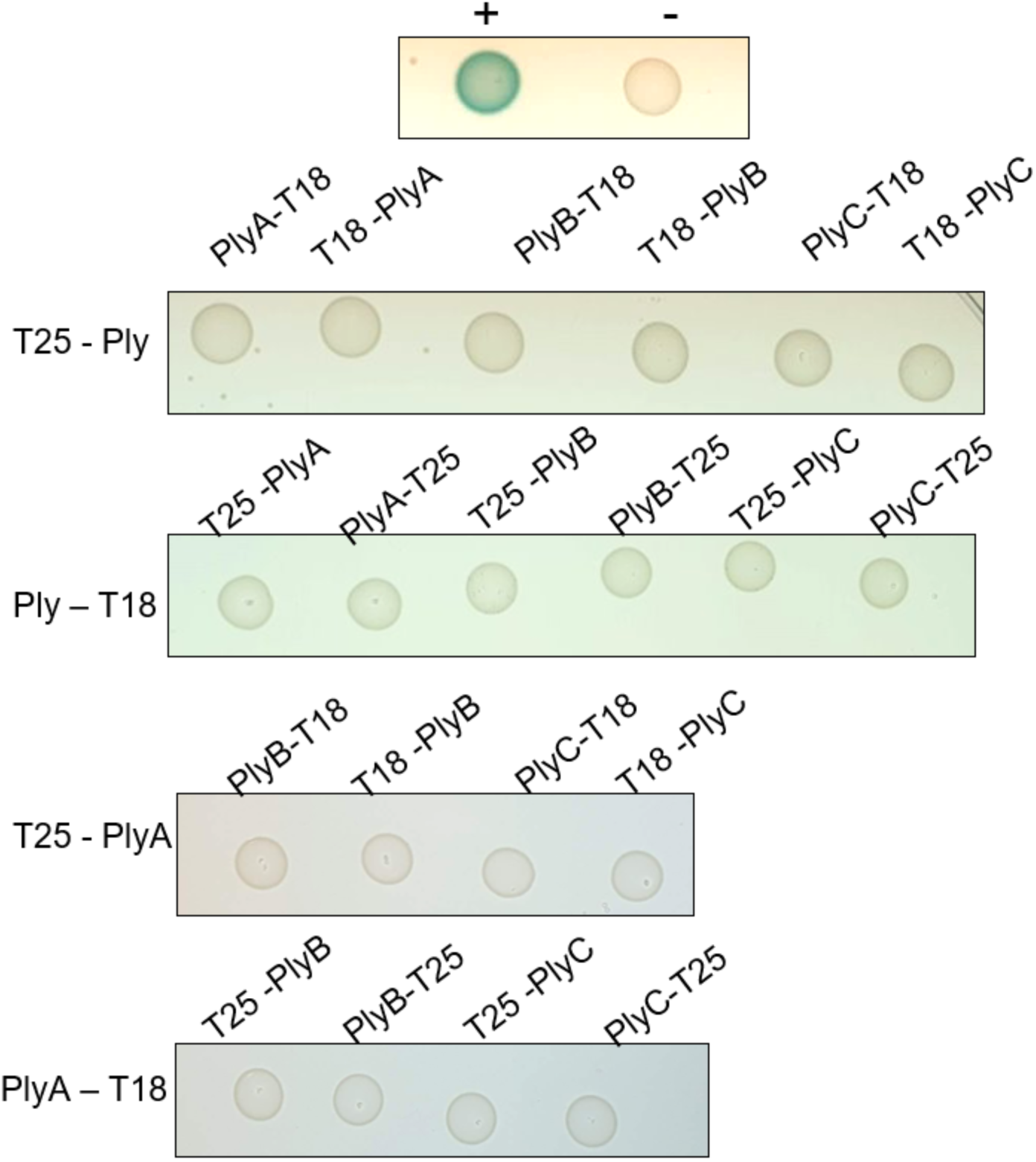
Bacterial two-hybrid analysis. Ply does not interact with any of the other proteins encoded on the operon. PlyA does not interact with PlyB and PlyC. Ply was tagged with the T25 subunit at the N-terminus and the T18 subunit at the C-terminus. PlyA, PlyB and PlyC were tagged with both the T25 and T18 subunits at the N- and C- termini. Representative of three biological replicates. pKT25 and pUT18 plasmids were co-transformed as negative control (−) and pKT25 – zip and pUT18C – zip as a positive control (+).

**Fig. S3.**
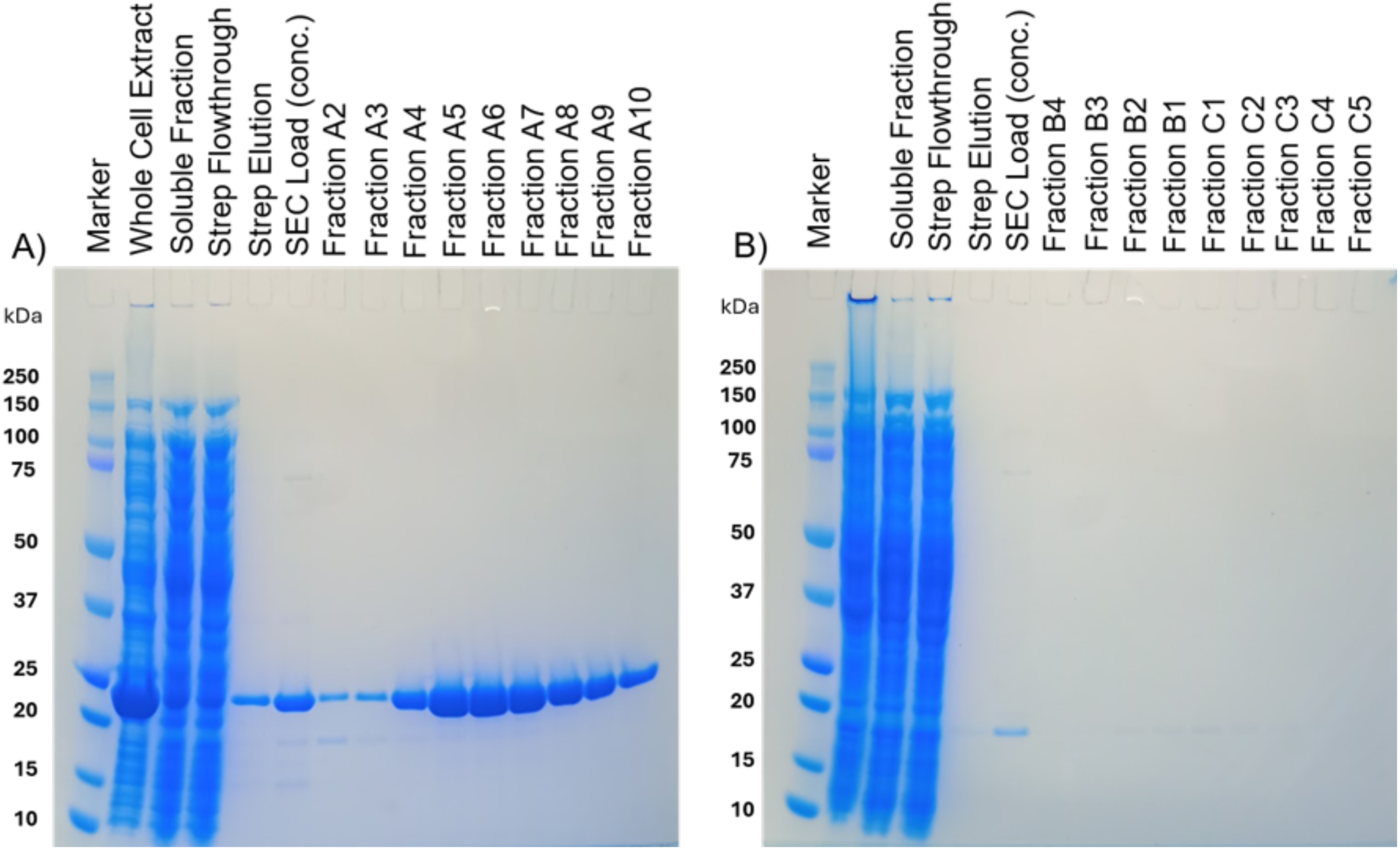
Purification of PlyB and PlyC proteins. SDS-PAGE loaded with protein fractions resulting from the purification of PlyB (A) and PlyC (B) using by affinity chromatography (Streptactin-XT) followed by size-exclusion chromatography (Superdex S75).

